# DNA replication dynamics are associated with genome composition in *Plasmodium* species

**DOI:** 10.1101/2024.09.18.613472

**Authors:** Francis Isidore Garcia Totañes, Sarah E. Chapman, Subash Kumar Rai, Mathew J. K. Jones, Michael A. Boemo, Catherine J. Merrick

**Author notes:** Address correspondence to: Catherine J. Merrick or Michael A. Boemo, Department of Pathology, Cambridge University, Tennis Court Road, Cambridge, CB2 1QP, UK.

## Abstract

*Plasmodium* species have variable genome compositions: many have an A/T-content of at least 80% while others are similar in composition to human cells. Here, we made a direct comparison of DNA replication dynamics in two *Plasmodium* species whose genomes differ by ∼20% A/T-content. This yielded fundamental insights into how DNA composition may affect replication. The highly A/T-biased genome of *P. falciparum* showed unusual replication dynamics that were not observed in the more balanced *P. knowlesi* – which had dynamics more like those of human cell lines. We observed that replication forks moved 50% slower in *P. falciparum* than in *P. knowlesi.* In *P. falciparum, r*eplication forks slowed down over the course of S-phase whereas in *P. knowlesi*, fork speed increased as in human cells. Furthermore, in both *P. knowlesi* and human cells, replication forks were strikingly slowed by sequences of particularly high A/T-bias, but in *P. falciparum*, although replication forks were inherently slow, they were not particularly slow in such biased sequences. Thus, the replisome of *P. falciparum* may have evolved alongside its extremely biased genome, making it unusually robust to sequence bias. Since several antimalarial drugs act to stall DNA replication, this study may have implications for the effectiveness of, and development of, antimalarial therapies.

## INTRODUCTION

*Plasmodium* parasites are the causative agents of malaria in humans and many other species. These early-diverging protozoa are unusual amongst eukaryotes in several respects. One of these is their unique form of cell division, called ‘schizogony’, which is syncytial rather than binary. Parasites inside erythrocytes – the pathogenic stage of malaria – divide asynchronously to produce multiple nuclei within a single cytoplasm followed by a single coordinated cytokinesis that produces dozens of daughter cells called merozoites. Cell division in other stages of this parasite’s complex lifecycle differ from schizogony but are also syncytial. Some resemble ‘endopolygeny’: repeated genome replications with neither karyokinesis nor cytokinesis until a final mass division stage (1). The process of schizogony has been much studied in recent years due to its implication in per-nucleus control of replication by non-diffusible regulators (conventional cyclin/CDK biology is lacking in *Plasmodium*), cell-cycle timing, and the control of nuclear/cytoplasmic ratios (2–4).

A striking feature of *Plasmodium* species is their highly variable genome composition. The six *Plasmodium* species that infect humans have all been sequenced, as have dozens of other non-human malaria species (5). The genomes are generally small (< 25 Mb) and largely orthologous, except in their large families of species-specific virulence genes. Many *Plasmodium* genomes, however, have an A/T-content over 80%, while others are much more balanced. For example, the most severe agent of human malaria, *P. falciparum,* has 80.7% A/T-content whereas the second most important parasite, *P. vivax,* has only ∼58% A/T-content (6).

The evolutionary reason for this bias is unknown, although it has been posited that a high rate of G/C → A/T substitutions maintains the bias (7). We hypothesised that the dynamics of DNA replication might be heavily affected by such a dramatically biased genome composition and that two related parasites whose genomes differ widely in A/T-content should exhibit differences in DNA replication fork speed. We therefore set out to compare genome replication dynamics in the only two human malaria species that are amenable to culture: *P. falciparum* (80.7% A/T) and *P. knowlesi* (61.4% A/T) (5).

We have previously compared schizogony in these two species at the cellular level (2), showing that they have both similar and divergent features. The *P. knowlesi* cell cycle was almost twice as fast as *P. falciparum* (∼30 versus 48h) and fewer merozoites were made per schizont. However, the parameters of schizogony were relatively similar: active replication (S-phase) took up a similar portion of the whole cell cycle, nuclei replicated asynchronously, and 20% to 100% of existing nuclei at any stage could replicate at once. We also followed the dynamics of origin recognition complex (ORC) in *P. falciparum* (8) and showed that this does not determine active replication: all nuclei throughout S-phase contained ORC, which was synthesised prior to the onset of S-phase. Rather than ORC being degraded or shuttled for each round of genome replication, PCNA may be the ‘limiting factor’ for active replication in *P. falciparum* since PCNA1 was recently shown to accumulate exclusively in actively replicating nuclei (3).

Subsequently, we examined *P. falciparum* replication at the molecular level, using both ORC-ChIP and the state-of-the-art DNAscent algorithm (9), which can determine sites of origin firing and speed of replication forks across the genome (8). We found that origins were placed very densely throughout the *P. falciparum* genome, without sequence-specificity beyond a preference for relatively high G/C content (in a very low-G/C genome). We found that replication forks moved slowly (2,8,10), were prone to stalling (10), and were slower towards the end of schizogony than the beginning (8,10), in contrast to the normal trend seen in human cells undergoing binary fission (11,12). Finally, we found that replication forks moved much faster in transcriptionally silent genes, suggesting that replication/transcription conflicts were particularly problematic for efficient replication in this genome (8).

Here, we measured DNA replication dynamics in *P. knowlesi* for the first time. Its genome, unlike *P. falciparum*, has an A/T-content of ∼60% which is similar to that of human cells. This revealed key differences that inform our general understanding of DNA replication dynamics. Forks moved ∼50% faster in *P. knowlesi* than in *P. falciparum* but were impeded by highly A/T-biased sequences. We observed the same trend in replicating human DNA. In sharp contrast, *P. falciparum* appears to have evolved a replisome that moves relatively smoothly through a highly A/T-biased genome, but nevertheless suffers from excessive stalling and is particularly impeded by active transcription.

## METHODS

### Parasites

Continuous *P. knowlesi* A1-H.1 cultures were grown in RPMI 1640 (Sigma, R4130) supplemented with 2.3 g/L sodium bicarbonate, an additional 2 g/L glucose (final concentration of 4 g/L), 0.05 g/L hypoxanthine (Sigma), 5 g/L Albumax II (Invitrogen), 10% horse serum (heat inactivated, Gibco) and 25 μg/mL gentamicin (Melford Laboratories), i.e. complete media, in 2% haematocrit O+ human red blood cells (NHS Blood and Transplant) in 3% oxygen, 5% CO_2_ and 92% nitrogen gas mixture at 37°C. To produce tightly synchronised parasites, schizonts were harvested using a 55% Nycodenz AG (ProteoGenix) gradient (v/v in RPMI) from 100% Nycodenz stock pH 7.5 (27.6% w/v Nycodenz, 5 mM Tris HCl, 3 mM KCl, 0.3 mM CaNa_2_·EDTA). Schizonts were incubated for 2 hours in complete media with either 1.5 μM compound 2 or 150 nM ML10 (LifeArc) (13). Compound 2 or ML10 was washed off with RPMI and mature schizonts were allowed to reinvade in 25% haematocrit red blood cells in complete media in the abovementioned gas mixture at 37°C for 1 hour at 300 rpm. The remaining schizonts were removed using 55% Nycodenz and the bottom layer was collected to produce a tightly synchronised culture (0 to 1 hours post invasion, referred to as 0 hpi).

### Generation of genetically modified parasites

To generate the donor DNA for *P. knowlesi* A1-H.1 *orc1* (PKA1H_130007800) C-terminal tagging, sequences approximately 800 bp upstream and 800 bp downstream from the stop codon, termed HR1 and HR2 respectively, were PCR-amplified with primers containing overhangs (with Tm of at least 56°C). The sequence containing 3xHA-T2A-NeoR was amplified from the pSLI-ORC1-3xHA plasmid from our previous *P. falciparum* work (8). The three fragments (HR1, 3xHA-T2A-NeoR, HR2) were fused together as described as three-step nested PCR in Mohring et al., 2019 (14). The donor DNA was cloned into a pCR2.1 plasmid using a TA Cloning Kit (Invitrogen). pCas9/sg plasmid (courtesy of Robert W. Moon, (14)) was digested with BtgZ1 (NEB) for 2 hours and purified using QIAquick PCR Purification Kit (Qiagen). Guide RNA sequence was ordered as oligos (Tm = 51°C) flanked with adaptor sequences matching the ends of the digested pCas9/sg vector and annealed by heating to 95°C then slowly cooling down using a thermocycler. The final CRISPR-Cas9 plasmid was assembled using NEBuilder HiFi DNA Assembly Master Mix (NEB) with 100 ng of digested vector and a 1:2 vector to insert molar ratio. The resulting mixture was ethanol precipitated and was transformed into electrocompetent PMC103 cells. Plasmids with guide RNA and donor DNA inserts were sent for Sanger sequencing for confirmation. Final plasmid maps are shown in Supplementary Figure 1A and B, and primer sequences used are listed in Supplementary Table 1.

Schizonts from thymidine kinase-expressing *P. knowlesi* A1-H.1, courtesy of Julian Rayner’s lab (2), were purified using 55% Nycodenz and were allowed to mature in complete media with 1.5 μM Compound 2 (LifeArc) (13) for 3 hours prior to transfection. Compound 2 was washed off and 20 μL of schizonts were allowed to recover in complete media at 37°C for 20 minutes. The schizonts were transfected with 20 μg of the final CRISPR-Cas9 plasmid and 60 μg of the donor DNA (circular plasmid) using an Amaxa 4D-Nucleofector and a P3 Primary Cell 4D-Nucleofector Kit (Lonza). The transfected schizonts were then transferred into a 1.5 mL microcentrifuge tube with 500 μL of complete media and 200 μL of red blood cells. The tube was incubated at 37°C in a thermomixer set at 550 rpm for 30 minutes then the contents were transferred into a 100 mm tissue culture dish with 4.5 mL of prewarmed complete media and cultured as usual. Once the selected parasite line reached about 1% parasitaemia, parasites with successful integration were then selected with 400 μg/mL of G418 (Merck).

Successful transfection, gene tagging and the absence of wildtype parasites were confirmed by PCR (Supplementary Figure 1C) and gene tagging was confirmed by western blot of fractionated protein lysates (Supplementary Figure 1D). Crude parasite protein lysates from tightly synchronised parasites were fractionated into cytosolic, nuclear soluble and nuclear insoluble fractions as previously described in Voss *et al.* 2002 (15). Briefly, saponin-lysed parasites were incubated for 5 minutes in ice-cold lysis buffer containing 20 mM HEPES, pH 7.8, 10 mM KCl, 1 mM EDTA, 1 mM DTT, 1 mM PMSF, 0.65% Nonidet P-40. Cytoplasmic protein fraction (supernatant) was obtained by centrifugation at 2500 x g for 5 minutes. The pellet was then incubated in extraction buffer containing 20 mM HEPES, pH 7.8, 800 mM KCl, 1 mM EDTA, 1 mM DTT, 1 mM PMSF, 1x Pierce protease inhibitor (Thermo Fisher Scientific) shaking at 2000 rpm at 4°C for 30 minutes. Centrifugation was done at 13000 x g for 30 minutes to separate the soluble and insoluble nuclear proteins (supernatant and pellet, respectively). Western blot of the fractionated samples was probed using rat anti-HA (Roche, clone 3F10) antibodies diluted 1:1000 in 2% milk. As control, the western blot membrane was also probed using mouse monoclonal anti-*P. falciparum* GAPDH antibody obtained from The European Malaria Reagent Repository (clone 13.3), and rabbit polyclonal anti-histone H4 antibody (Abcam, ab10158). SuperSignal West Pico PLUS Chemiluminescent Substrate (Thermo Fisher Scientific) was used for detection. Visualisation and imaging were done using an Azure Biosystems 500Q imager. The resulting modified parasite line, i.e. *P. knowlesi* ORC1-3xHA + pTK-Pyr, was utilised for the subsequent immunofluorescence and nanopore sequencing experiments.

### Immunofluorescence

Tightly synchronised *P. knowlesi* ORC1-3xHA + pTK-Pyr culture was incubated in 100 μM BrdU (Sigma) 30 minutes prior to collection of samples for immunofluorescence staining. Parasite smears were made every 2 hours from 14 hpi to 32 hpi. Thick smears were fixed with 4% formaldehyde in PBS for 10 minutes followed by permeabilisation in 0.2% Triton-X100 for 15 minutes. Slides were incubated in 1U/mL of DNAse I (Thermo Fisher Scientific) in 1X DNAse reaction buffer with MgCl_2_ (Thermo Fisher Scientific) for 45 minutes in a humidified chamber. DNAse solution was rinsed off with PBS and slides were blocked in 1% BSA (Sigma) with 0.1% Tween-20 (i.e., block) for 1 hour. Primary antibody labelling was done for 1 hour using mouse anti-BrdU (Cytiva, clone BU-1) and rat anti-HA (Roche, clone 3F10) antibodies diluted 1:500 in block. Three 5-minute washes with block were done prior to incubation in secondary antibodies (Thermo Fisher Scientific Alexafluor goat anti-mouse 488 and Alexafluor goat anti-rat 594) diluted at 1:1000 in block for 1 hour. Three 5-minute final washes were done with block, with the second wash replaced with DAPI (Thermo Fisher Scientific) at 2 μg/mL in PBS. All incubation steps were done at room temperature. Slides were allowed to cure overnight at room temperature using ProLong Diamond Antifade Mountant (Invitrogen) and were stored at 4°C prior to visualisation.

Images were acquired using a Nikon Microphot SA microscope with a Qimaging Retiga R6 camera at 1000x magnification. ImageJ (16) was used to convert all raw images to 32-bit and to identify and create the corresponding regions of interest (ROIs) on nuclear signal in the DAPI channel images. Without creating any further image adjustments, the ROIs were determined using ImageJ Threshold set to MaxEntropy (17). These ROIs where then used to measure the integrated signal density in all images taken using the different fluorescent channels. The resulting data were analysed and plotted using GraphPad Prism v9.3.1. For the representative images, fluorescence signal brightness was minimally adjusted to show contrast between background and actual signal. Representative images were pseudo-coloured and merged using ImageJ.

### Datasets for identification of active replication forks and origins

To produce data sets for the identification of active replication forks and origins, tightly synchronised parasites were incubated in 20 μM 5-ethynyl-2-deoxyuridine (EdU, Thermo Fisher Scientific) for 7.5 minutes, then 200 μM 5-bromo-2-deoxyuridine (BrdU, Merck) for another 7.5 minutes, followed by 2 mM thymidine for 1 hour and 45 minutes prior to parasite harvest. Nascent DNA labelling was done at 20 and 23 hpi, representing early and mid-schizogony, respectively. High molecular weight parasite genomic DNA was extracted using a MagAttract High Molecular Weight Genomic DNA kit (Qiagen) or Blood & Cell Culture DNA Mini Kit with Genomic Tip 20/G (Qiagen) and stored at 4°C. Long DNA fragments were enriched using SRE-XL short read eliminator buffer (PacBio) prior to library preparation and nanopore sequencing. DNA barcoding was done on 1 μg of high molecular weight genomic DNA using Oxford Nanopore ligation sequencing kit (SQK-LSK109) and barcode expansion kit (EXP-NBD104). Sequencing was done using MinKNOW software version 23.04.5 set to produce fast5 output files with a Nanopore MinION device and R9.4.1 flow cells for 72 hours or until the pores in the flow cell were fully depleted.

### Data analysis

To call replication forks and origins on single molecules, Oxford Nanopore sequencing reads were basecalled and demultiplexed with Guppy (v5.0.11) and aligned to the *P. knowlesi* PkA1H1_v1 assembly (https://www.ncbi.nlm.nih.gov/datasets/genome/GCA_900162085.1/) with minimap2 (v2.17-r941). Only sequences with an alignment length ≥ 10 kb and mapping quality ≥ 20 were analysed. The probability of BrdU and EdU at thymidine positions along each read was assigned by DNAscent detect (v3.1.2), and these probabilities were parsed into replication fork and origin calls by DNAscent forkSense. Bedgraphs of base analogue calls on single molecules were generated using the dnascent2bedgraph utility. *P. falciparum* DNAscent data were obtained from our previously published data (8). Human DNAscent data were obtained from RPE1 cells incubated in 50 μM of EdU for 5 minutes, then labelled with 50 μM of BrdU for 10 minutes and finally 100 μM of thymidine for 20 minutes with 3 times PBS wash in between addition of nucleotides (18).

Available data sets for both *P. falciparum* and *P. knowlesi* (i.e., coding sequences, low complexity regions, tandem repeats, *var* and *SICAvar* gene data, as well as *P. knowlesi* Strain H microarray transcriptomics data (19), and HP1 log2ratios (20)) were downloaded from PlasmoDB (5). Coordinates of *P. knowlesi* data sets that were mapped to Strain H genome assembly (GCA_000006355.3) were lifted over to the A1-H.1 assembly (GCA_900162085.1) using a chainfile generated through flo software version 1.1.0 (21). Interstitial telomere repeat sequence (ITS) density in *P. knowlesi* A1-H.1 (v66) genome and *P. falciparum* 3D7 (v66) was calculated by counting the occurrences of GGGTTYA sequences in both forward and reverse strand per 100 bp region using Python (20). Similarly, the density of human ITS was calculated by counting the occurrence of TAGGGAGG and TTATGGAGG sequences (22) per 100 bp in the human genome (GRCh38.p13). Visualisation of chromosome features was done using kartoploteR (23).

BEDtools v2.30.0 (24) was used for file format conversions, comparison of multiple data sets, and identifying genes overlapping with areas of interest in the genome. Matrix plots (computeMatrix) were done using deepTools 3.5.0. (25). G/C-content of DNAscent origins was calculated by obtaining the sequence from 49 bases upstream to 50 bases downstream of the given origin midpoint. G/C-content of forks was calculated by looking at the last 100 bp of the moving fork (i.e. first 100 bases of the coordinates of a left moving fork or the last 100 bases of a right moving fork). Fork speed calculation was done on all forks identified by DNAscent that were not located at the nanopore read ends, or joined at the replication origin or termination site. Fork speed was calculated by dividing the distance (in kb) that the fork travelled during the EdU and BrdU pulses by 15 minutes (the combined duration of the two pulses) giving the fork speed in kb/min. Fork speed z-score was calculated by dividing the difference between a given fork speed and the mean fork speed by the overall standard deviation. This allows clearer visualisation of deviations of fork speed from the mean as replication forks pass through any given sequence. To determine any effect of A/T-content on fork speed, a given genome was divided into 100 bp bins and sequences with “N” were removed. The bins were then divided by into 10 clusters, grouped by G/C-content (i.e. from 0 ≤ A/T% ≤ 10 to 90 < A/T% ≤ 100). Only clusters with > 50 bins and bins with fork speed data were included in the analysis, and p-value was corrected for multiple comparisons (p-value ≤ 0.001 were considered significant). Mean z-score ± standard error over genomic regions was calculated and plotted using deepTools.

Scatterplots were generated using GraphPad Prism v9.3.1. Test for normality, Mann-Whitney U test, and t-test (two-tail) were done using SciPy (26). Hypergeometric p-value for the comparison of origin G/C-content against the whole genome was calculated by obtaining random 100 bp sequences from the genome with the same quantity and chromosomal distribution as the origins. The mean G/C-content of the group of random sequences was then computed. One million randomisations were done to obtain random G/C values against which the actual G/C-content was compared.

Human, *P. knowlesi*, and *P. falciparum* polymerase (α, δ, and ε) and helicase (RECQ1 and WRN) protein sequences were downloaded from Uniprot (27) and sequence alignments were done using Clustal Omega Multiple Sequence Alignment (28). To predict protein-DNA complex structures, 500 base DNA sequences with varying G/C-content (i.e. 80%, 40% and 20%) were obtained from chromosome 1 of the human genome (GRCh38.p13), *P. knowlesi* A1-H.1 (v66) genome and *P. falciparum* 3D7 (v66), respectively. These were then analysed together with the polymerase/helicase sequences using Alphafold 3 (29).

## RESULTS

### Comparison of genome characteristics in *P. knowlesi* versus *P. falciparum*

Unlike the *P. falciparum* genome, which has an overall A/T-content of 80.7%, the *P. knowlesi* genome has an A/T-content of 61.4%, very close to that of the human genome (59.1% (30)). The median A/T-content of coding sequences in *P. knowlesi* (59.7%) is similar to that of the whole *P. knowlesi* genome, whereas *P. falciparum* coding sequences exhibit an A/T-content (75.8%) that is markedly reduced relative to the whole *P. falciparum* genome (albeit still unusually high for coding sequences). Table 1 summarises the genomic features in both species. The nucleotide composition of the whole genome and other genomic features in both species are shown in Figure 1.

**Figure 1.**
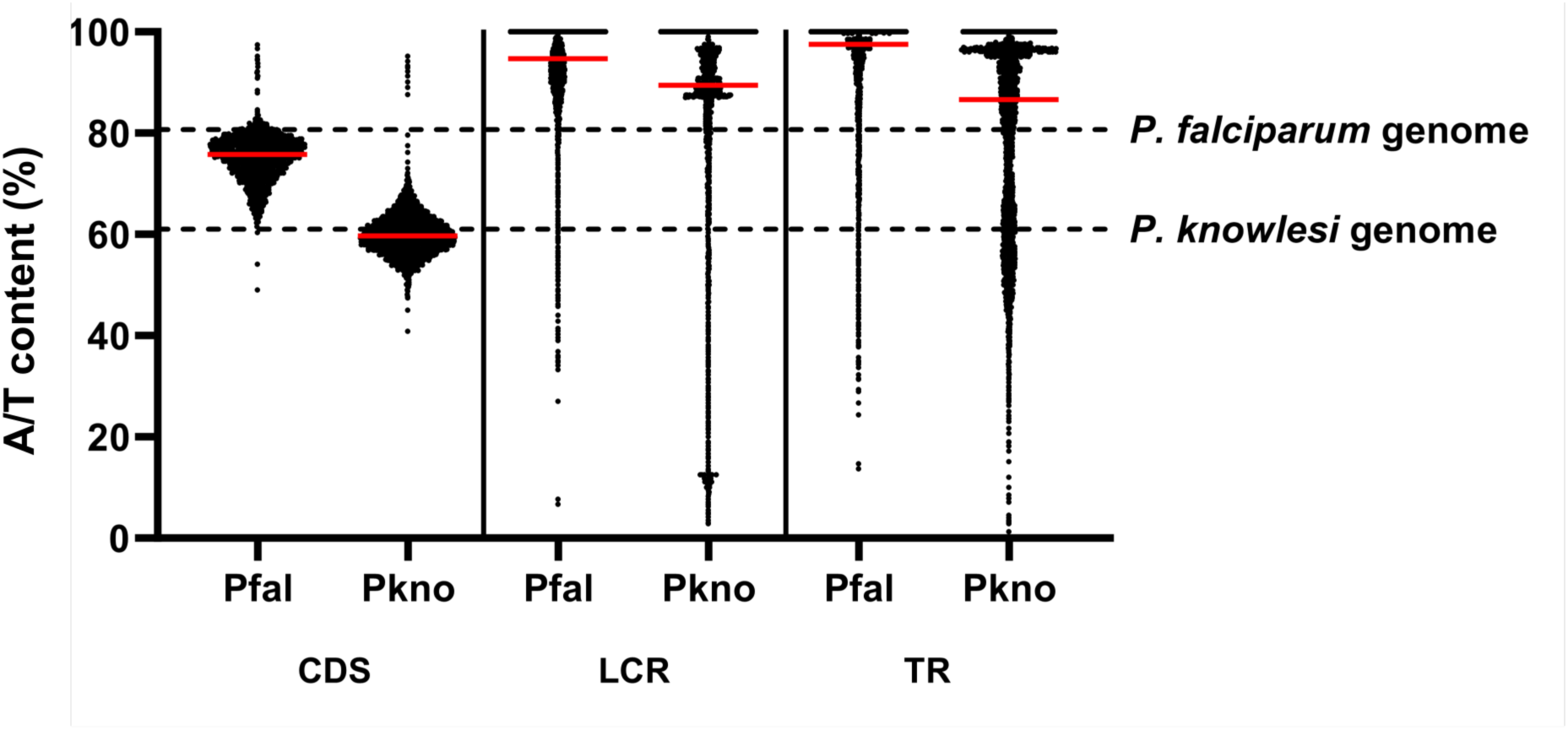
Comparison between the A/T content in coding sequences (CDS), low complexity regions (LCR), and tandem repeats (TR) in *P. falciparum* and *P. knowlesi*. The overall A/T contents of the two species are represented by the dashed line while red bars represent the median A/T content.

**Table 1.** Comparison of genome size, A/T-content, and other genomic features between *P. falciparum* and *P. knowlesi*. CDS, coding sequence; LCR, low complexity region; TR, tandem repeat.

These two *Plasmodium* species are orthologous in most of their genes, with 3803 orthologous gene pairs (31) but they differ in their virulence gene families – particularly in the large gene families that encode immunodominant antigens expressed on the surface of infected erythrocytes. There are 214 genes that belong to the schizont-infected cell agglutination variant antigen (*SICAvar*) family in *P. knowlesi*. These genes are distributed throughout the genome, mainly overlapping with interstitial telomere repeat sequence (ITS) and heterochromatin protein 1 (HP1) enriched regions (Figure 2A). Similarly, the 64 genes in the *var* gene family of *P. falciparum* overlap with HP1-rich regions in the genome; however, a majority of these genes are located in telomeric and subtelomeric regions (Figure 2B) where they are silenced by conventional telomere-associated heterochromatin.

**Figure 2.**
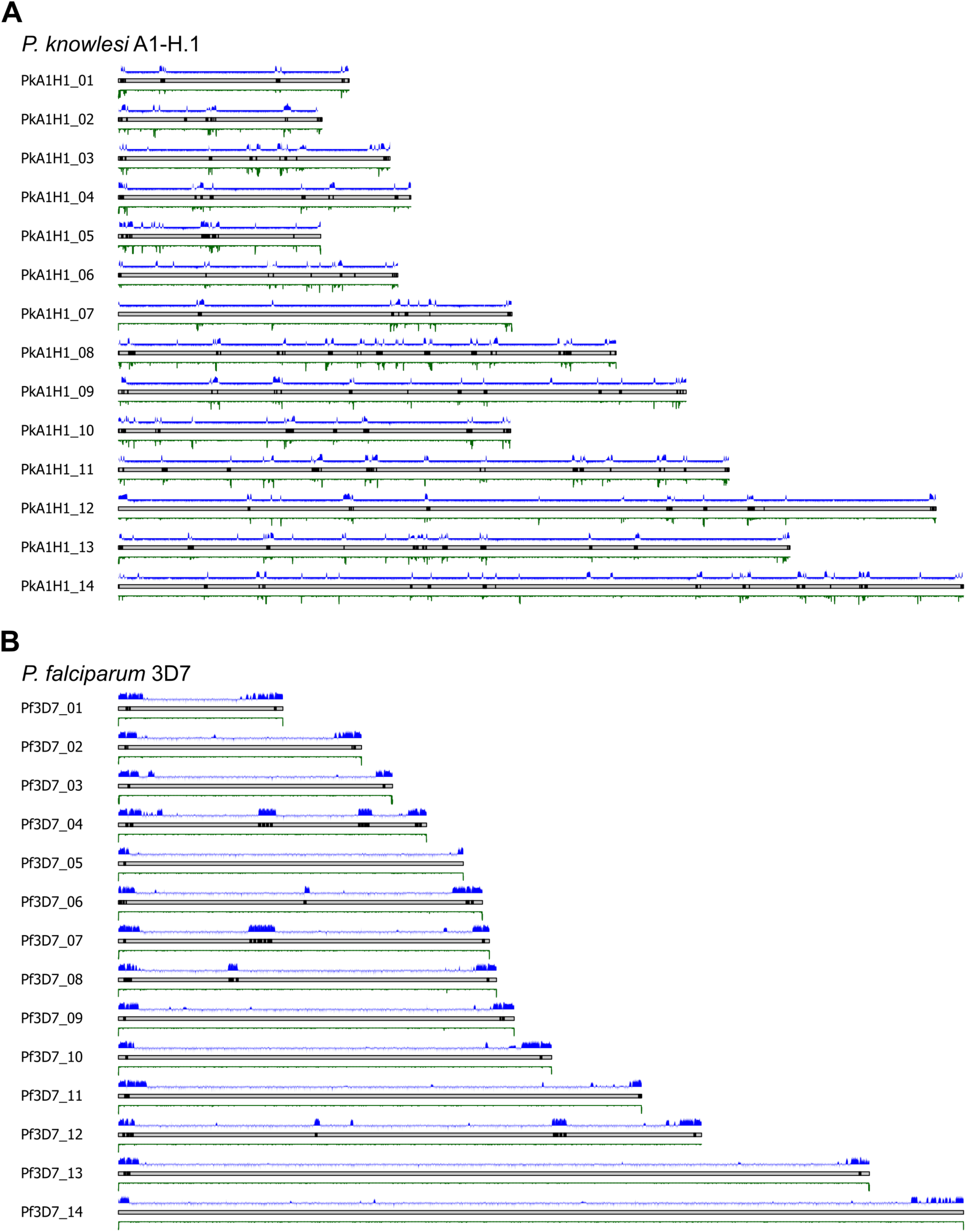
Chromosomal distribution of *var* or *SICAvar* genes (black bars), HP1 distribution (20) (log2ratio, blue), and ITS density (green) in (A) *P. knowlesi* A1-H.1 and (B) *P. falciparum*.

### The origin recognition complex shows similar cell-cycle dynamics in *P. knowlesi* and *P. falciparum*

To investigate the dynamics of DNA replication during blood-stage schizogony in *P. knowlesi*, we first used immunofluorescence to follow the dynamics of the origin recognition complex, ORC, in comparison with the dynamics of nascent DNA replication. We had previously performed analogous experiments in *P. falciparum* (8).

*P. knowlesi orc1* (PKA1H_130007800) was C-terminally tagged in thymidine kinase-expressing parasites that are capable of incorporating thymidine analogues such as BrdU and EdU during active DNA replication (2). Fractionation of crude parasite lysate showed that ORC1 could be detected in the cytosolic, soluble and insoluble nuclear fractions but the strongest signal was observed in the soluble nuclear fraction. This was consistent with the appearance of ORC1 in the immunofluorescence assays shown in Figure 3. For this experiment, tightly synchronised parasites were allowed to incorporate BrdU for 30 minutes prior to sample collection every 2 hours from 14 to 32 hpi, spanning the entire S-phase period (parasite cultures were monitored and samples were collected until a majority (> 60% of parasites) exhibited the typical fan-shaped schizont form). We first observed ORC1 at 16 hpi, about four hours prior to the first visualisation of active DNA replication. This was similar to what was previously observed in *P. falciparum,* where ORC1 was expressed before active DNA replication commenced (8). Similarly, ORC1 was observed in all nuclei from 20 to 32 hpi regardless of whether active replication was occurring or not. Parasite images taken at 24 and 26 hpi best exemplify this, where all nuclei have ORC1 signal while only one is actively replicating (Figure 3).

**Figure 3.**
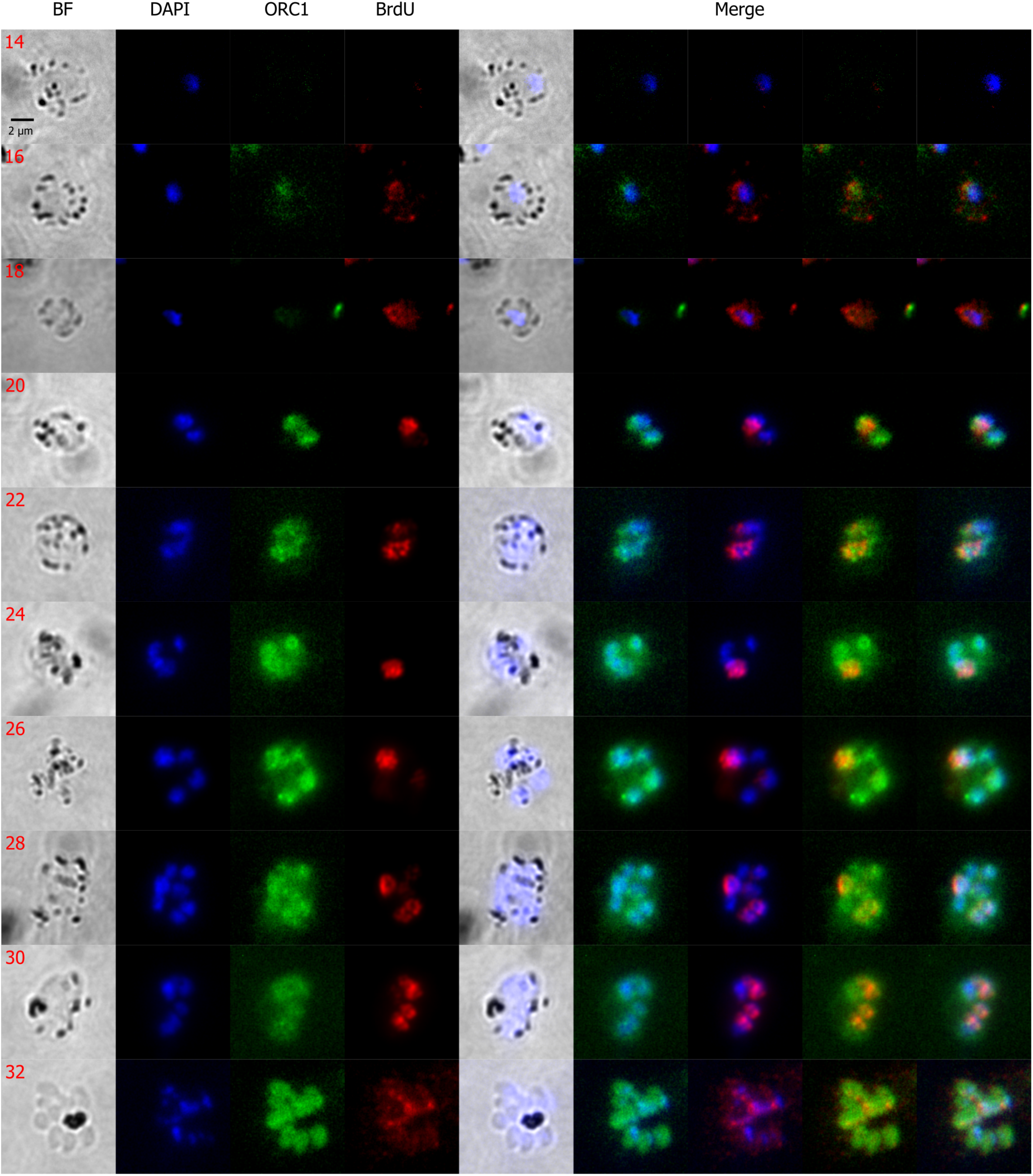
Representative immunofluorescence images showing ORC1 distribution and BrdU labelling in *P. knowlesi.* Synchronised *P. knowlesi* ORC1-3xHA + pTK-Pyr was incubated in BrdU for 30 mins at 2h intervals from 14 to 32 hpi. ORC1 (in green) and BrdU (in red) were probed using anti-HA and anti-BrdU antibodies, respectively. DNA was stained with DAPI (blue). Scale bar (2 μm) applies to all images.

Using ImageJ, fluorescence signals were quantified by measuring the integrated density of each signal (i.e. DAPI, ORC1 and BrdU) within the region of the parasite nuclei (Figure 4). As expected, the amount of DNA and the amount of ORC1 both increased by ∼10-fold as multiple nuclei were produced across the course of schizogony, while the amount of active DNA replication increased ∼7-fold. Interestingly, towards the end of schizogony ORC1 abundance was observed to peak at 28 hpi followed by a decrease at 30 and a slight increase at 32 hpi (Figure 4A). This pattern followed the trend of *orc1* gene expression that was previously reported in microarray data from *in vitro P. knowlesi* cultures (Figure 4D) (19). BrdU signal also reached the maximum median value at 28 hpi, before decreasing (Figure 4B). A similar trend was also observed in our previous timecourse experiments in thymidine kinase-expressing *P. knowlesi* that were allowed to incorporate EdU (2). Aside, however, from this reproducible pattern of fluctuation seen in late schizogony, the overall dynamics of ORC1 production and DNA replication in *P. knowlesi* were broadly similar to the pattern we previously reported in *P. falciparum*.

**Figure 4.**
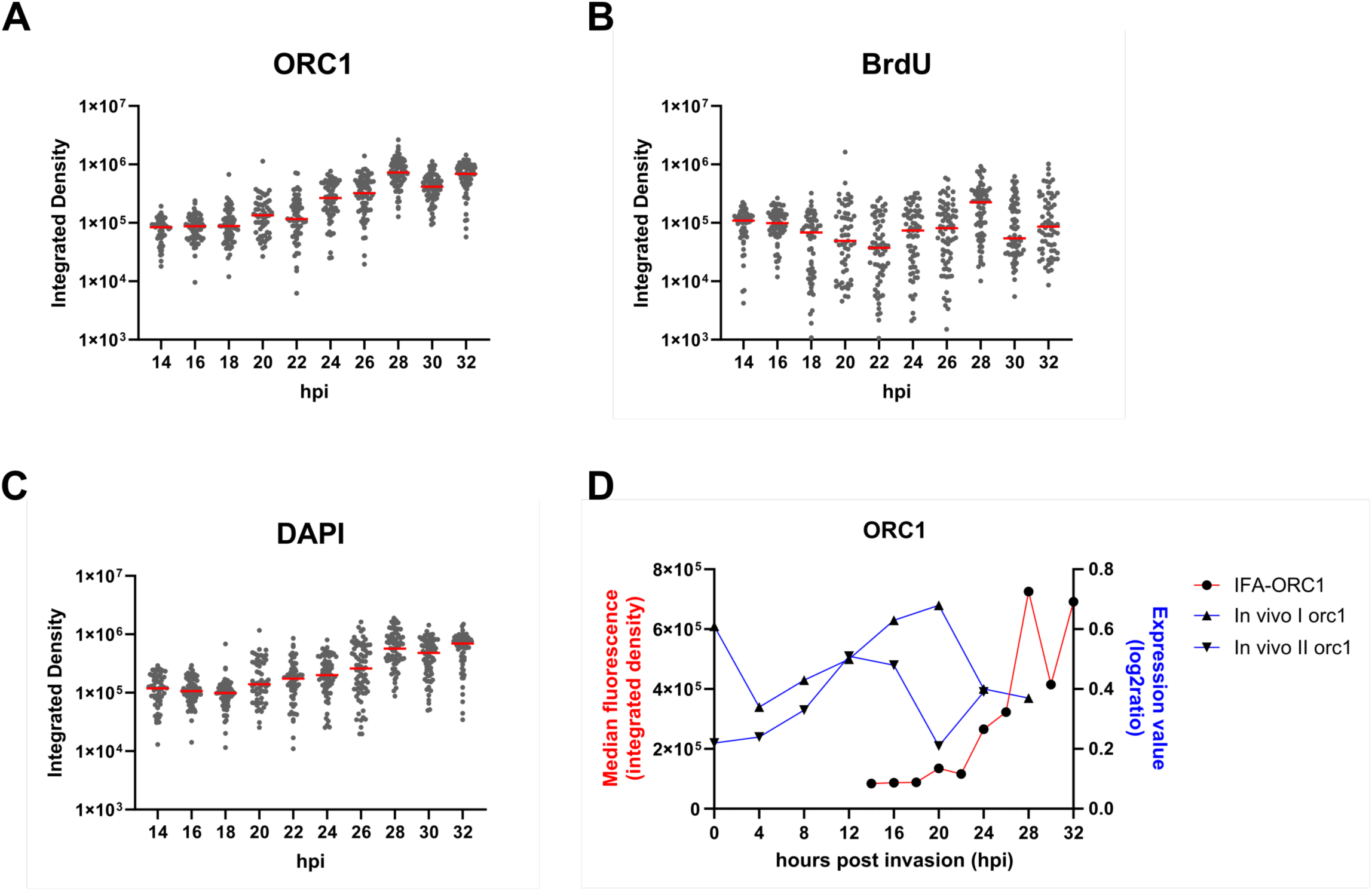
Quantification of immunofluorescence data shown in Figure 3. A) ORC1, (B) BrdU and (C) DAPI in tightly synchronised *P. knowlesi* ORC1-3xHA + pTK-Pyr from 14 to 32 hpi. Y-axes are shown in log scale to show differences in median integrated densities between time points (the same data for ORC1 are shown on a non-log scale in (D)). The median values are represented by red bars in the dot plots (n = 53-72 cells per timepoint). (D) Median ORC1 integrated density in comparison with published data from 2 replicates of *orc1* gene expression, measured in log2ratio from *in vivo* culture (19).

### Replication origins and replication fork dynamics differ in *P. knowlesi* compared to *P. falciparum*

Having defined the dynamics of ORC1 synthesis and nascent DNA replication at the cellular level in *P. knowlesi*, we moved on to examine replication at the single-molecule level. We hypothesised that genome composition might strongly affect replication dynamics, leading to a superficially similar cellular phenotype (i.e. schizogony) being generated by very distinct molecular-level patterns in *P. knowlesi* versus *P. falciparum*, the species we had studied previously (8).

High molecular weight genomic DNA was extracted from 6 biological replicates of tightly synchronised parasites labelled with EdU and BrdU at 20 and 23 hpi. These timepoints represent the beginning and middle of schizogony. DNA was barcoded, sequenced on the Oxford Nanopore Technologies platform, and analysed for active replication forks and origins via the DNAscent algorithm, which calls the positions of thymidine analogues EdU and BrdU in each sequenced molecule (9). For each replicate, 2 barcoded samples (i.e., one for each timepoint) were combined into a single library which was sequenced using a MinION R9.4.1 flow cell. Supplementary table 2 summarises the total number of forks and origins called by DNAscent in all 6 replicates.

Replication fork speed was strikingly faster in *P. knowlesi* than in *P. falciparum* (2) (Figure 5A). This was consistent with the hypothesis that highly A/T-biased DNA, unique to *P. falciparum,* may present particular difficulties for efficient replication. There was also a difference in how replication fork speed changed across the course of schizogony. In *P. knowlesi,* at the beginning of schizogony, forks were marginally but significantly slower compared with those at mid-schizogony (median fork speeds of 0.829 kb/min and 0.852 kb/min, respectively; p-value: 0.0387). Conversely, fork speed at the beginning of schizogony in *P. falciparum* was marginally higher that the speed at mid-schizogony (median fork speed of 0.589 kb/min and 0.548 kb/min respectively, p-value: 3.24E-08).

**Figure 5.**
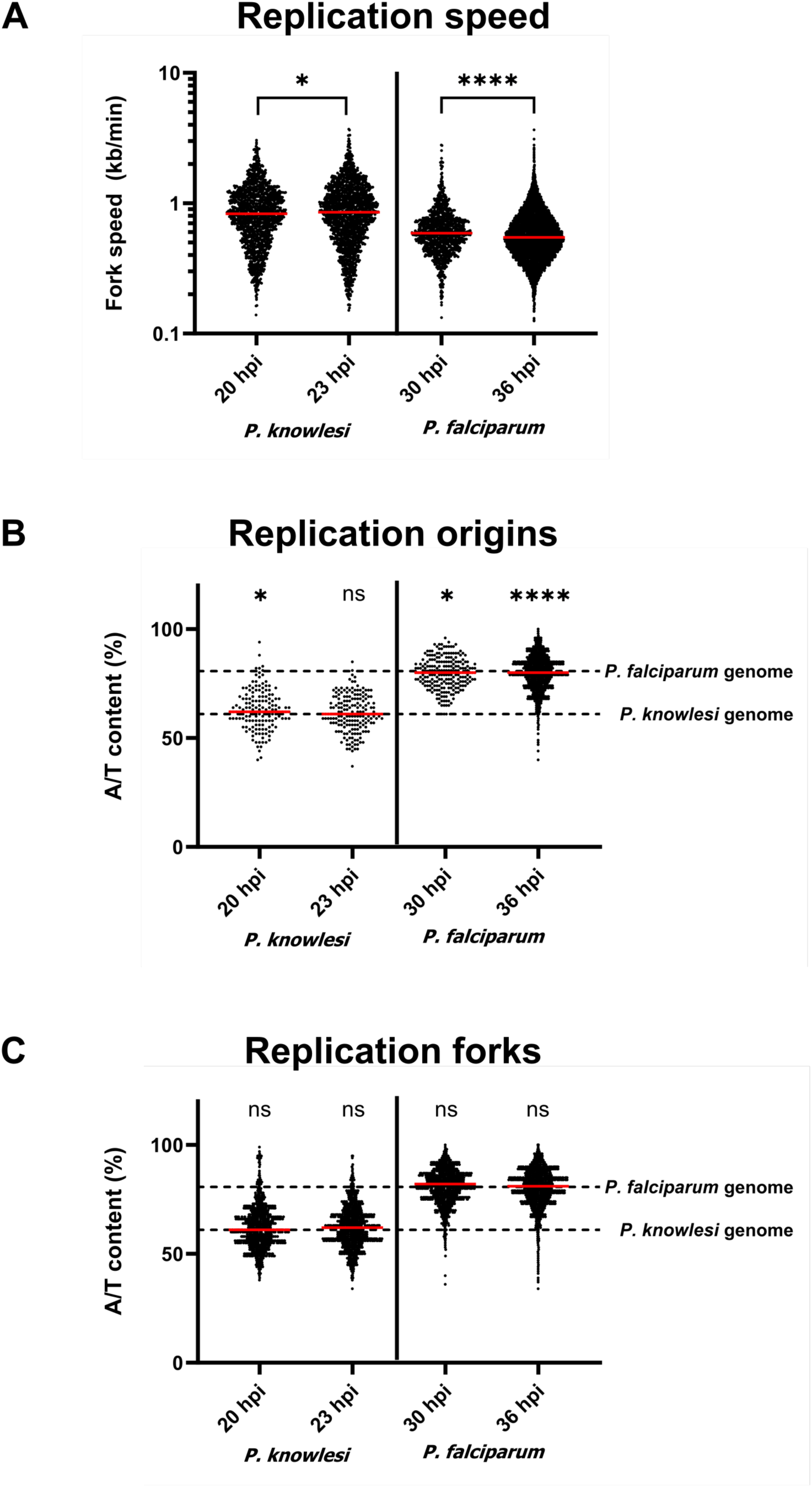
Replication fork and origin parameters in *P. knowlesi* versus *P. falciparum.* (A) *P. knowlesi* fork speed at 20 and 23 hpi in comparison with *P. falciparum* (at 30 and 36 hpi). A/T-content of (B) replication origins (100 bp at the centre of the origin), (C) forks (100 bp at the moving end of each fork). Hypergeometric p-values were calculated comparing origin and fork A/T content against the whole genome: * = 0.030, **** = 5.00E-06, ns = non-significant.

Concerning replication origins, those that fired at 20 hpi in *P. knowlesi* had a marginally but significantly higher A/T-content (median A/T: 63.0%) compared with the whole genome (hypergeometric p-value = 0.0431); however, there was no significant difference in origins that fired at 23hpi (median A/T: 62.0%, hypergeometric p-value = 0.290) (Figure 5B). By comparison, active origins at comparable timepoints, 30 and 36 hpi, in *P. falciparum* both had 80.0% A/T, slightly lower than the whole genome (hypergeometric p-values: 0.030 and 5.00E-06, respectively). (Of note, most of the active replication origins identified in *P. knowlesi* – 124 out of 154 (80.5%) at 20 hpi and 136 out of 145 (75.1%) at 23 hpi – were origins that fired during the analogue incorporation, while the rest were origins identified by taking the midpoint between two actively moving forks, i.e. left- and right-moving forks. The former method is likely to give the most accurate determination of origin placement.) As expected, no significant difference in A/T-content was seen between active forks and the whole genome for both species (Figure 5C).

We found no significant correlation between the distribution of *P. knowlesi* replication forks and origins and the placement of heterochromatin protein 1 (HP1) throughout the whole genome, similar to what we previously observed in *P. falciparum* (8) (Supplementary table 3 and Supplementary figure 2). However, the representation of forks did differ in virulence gene families – which are generally enriched in HP1. There was a significant enrichment of left- and right moving forks at the end and start of *var* genes, respectively, in *P. falciparum* (8) but this was not observed in the *SICAvar* genes of *P. knowlesi* (Supplementary figure 2).

### Replication fork speed is differently affected by active gene transcription in *P. knowlesi* versus *P. falciparum*

In our previous study of *P. falciparum,* we reported a striking bias in replication fork speed, with silent genes being traversed much faster than actively transcribed genes (8). This suggested that active transcription interfered greatly with efficient DNA replication in this genome. In *P. knowlesi,* we observed no clear relationship between active gene transcription and efficient replication fork movement (Figure 6).

**Figure 6.**
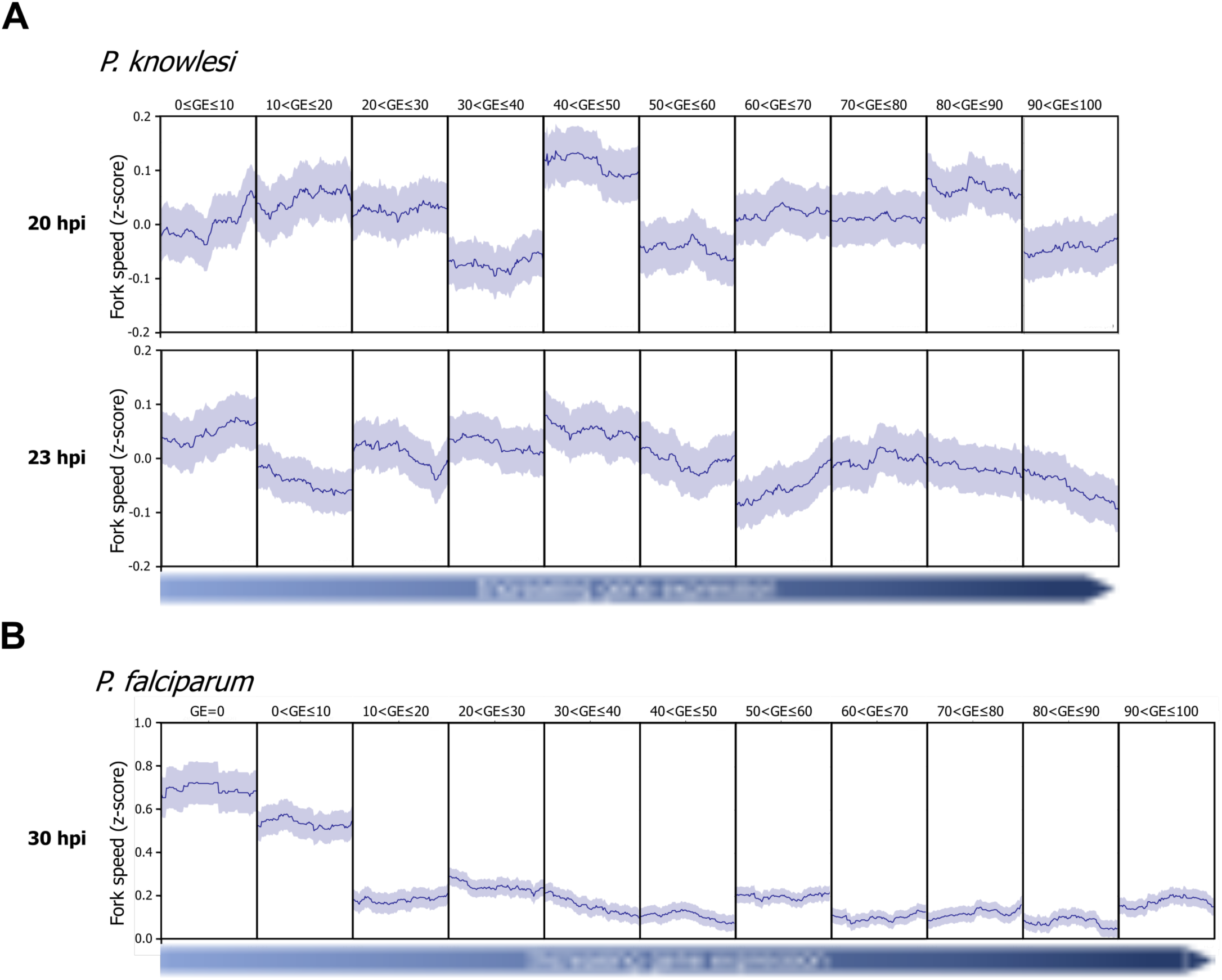
Comparison of replication fork speeds in silent and actively-transcribed genes in *P. knowlesi* versus *P. falciparum.* Mean fork speed z-score shown as dark blue line mapped along coding sequences of genes clustered by level of gene expression at the beginning and middle of schizogony in *P. knowlesi* (A) and *P. falciparum* (8) (B). The coding regions of genes within each cluster were scaled, with the left and right borders of the plot corresponding to the transcription start site and transcription end site, respectively. The standard error of the mean is shown as light blue shading above and below the mean fork speed.

### Replication fork speed is differently affected by genome composition in *P. knowlesi* versus *P. falciparum*

To determine whether genome composition can affect fork speed, the whole genome was divided into 100 bp bins and categorised for nucleotide content. These bins were divided into 10 clusters from 0 - <10%, to 90 - 100% A/T. Replication fork speed (mean z-score) over these sequence clusters was then calculated and plotted for *P. knowlesi* and *P. falciparum.* Since the results clearly differed, we then also analysed the same parameters in existing data sets from human cells, to determine which *Plasmodium* species represented the ‘norm’ versus the ‘exception’ (Figure 7).

**Figure 7.**
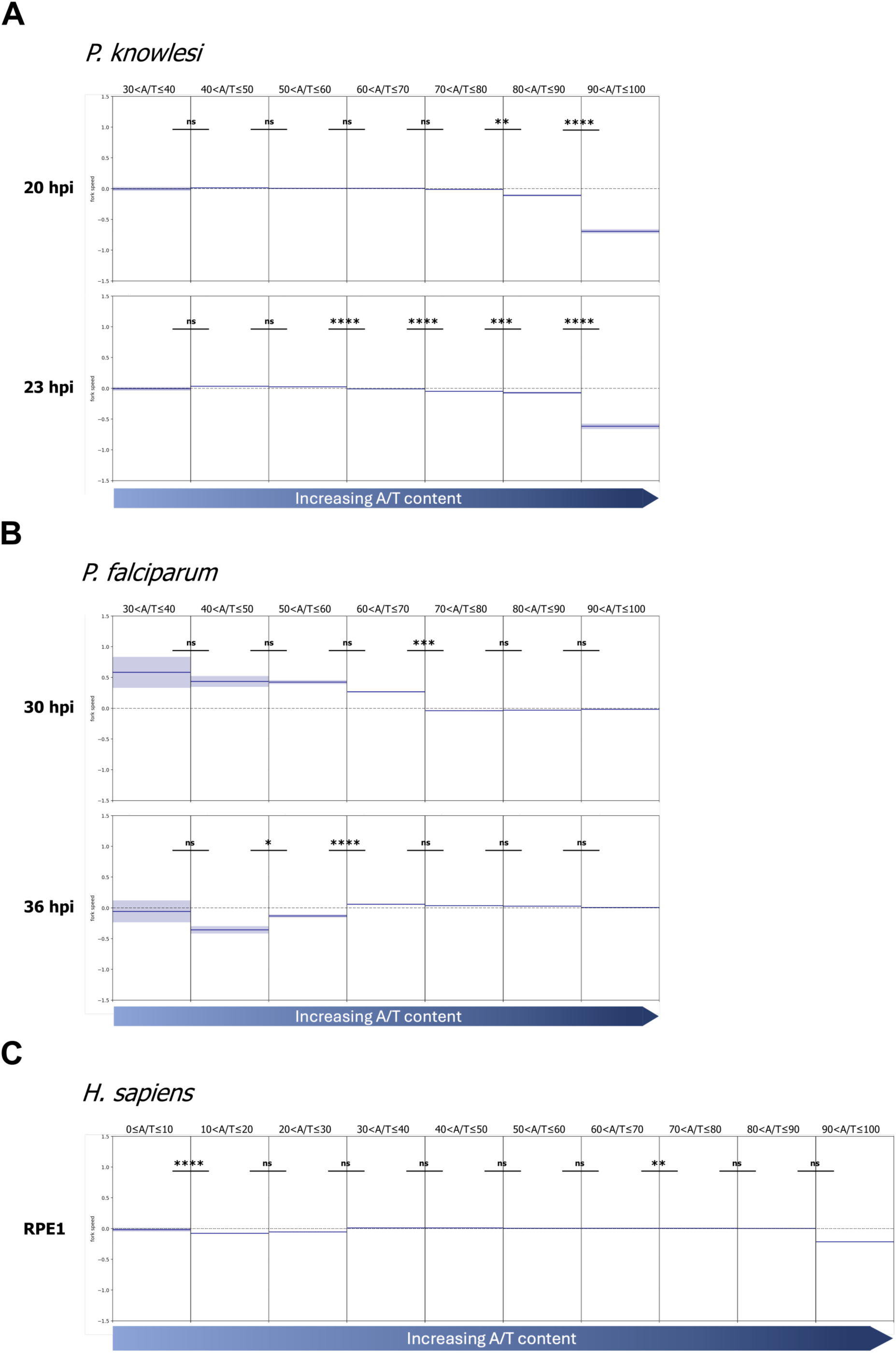
Comparison of replication fork speeds across regions with different A/T-content in *P. knowlesi*, *P. falciparum* and *H. sapiens.* Mean fork speed z-score ± standard error over genome sequences clustered by A/T-content. The whole genome of each species was divided into 100 bp regions (bins) and bins were clustered from 0 to <10% to 90 to 100% A/T; however, only clusters with > 50 bins were included in the analysis. The average fork speed z-score (representing positive or negative deviation from the overall mean fork speed, shown as dark blue lines) ± standard error (light blue shading) was calculated and Mann-Whitney U test was used to calculate significant difference between z-scores of neighbouring clusters. Dashed line marks 0 z-score, representing the genome-wide mean fork speed. p-value: * ≤ 1E-03, ** ≤ 1E-04, *** ≤ 1E-05, **** ≤ 1E-06

Replication forks in *P. knowlesi,* at both the beginning and middle of schizogony, were slowest in genomic regions with very high A/T-content (greater than 90%), compared to regions with A/T-content of 80 - 90% (p-values: 3.11E-14 and 6.17E-30, respectively). A similar trend was seen in data from human RPE1 cells: replication was strikingly slow in areas with A/T-content greater than 90% (p-value = 1.42E-32).

Outside this cluster, replication speeds were less variable, but fork speeds in regions of the *P. knowlesi* genome with A/T-content of 80 - 90% were still below the overall average fork speed at both early and mid-schizogony. In mid-schizogony, there was an increase in fork speed (above the overall average) in regions with relatively low A/T-content, i.e. 40 - 60%. Overall, replication of genomes with balanced nucleotide content – both *P. knowlesi* and *H. sapiens* – was most efficient in regions of ‘moderate to high’ G/C-content and was inefficient in highly A/T-biased regions.

On the other hand, in *P. falciparum*, there was no bias towards very slow replication in DNA with A/T-content above 90%: such regions replicated at the average speed for this species. There was, however, a significant increase in average fork speed in regions with A/T-content less than 70%, but this was seen only at the beginning of schizogony. At mid-schizogony, lower-A/T regions no longer favoured faster replication speeds, in fact there was a significant decrease in average fork speed observed at 36 hpi in replication forks traversing regions with 40 - 60% A/T-content.

To investigate possible reasons for the different trends in replication fork speeds amongst these species (Figure 7), we aligned protein sequences of the human, *P. knowlesi* and *P. falciparum* polymerases (α, δ, and ε), and of some representative secondary-structure helicases (RECQ1 and WRN), to identify similarities and differences. We hypothesised that the structure of *P. falciparum* replisome component(s) might be unusual. However, the polymerases and helicases in *P. knowlesi* and *P. falciparum* all had higher percentage identity than human and *P. knowlesi,* hence providing no insight into the greater similarity of fork movement dynamics of human and *P. knowlesi* (Supplementary table 4). Furthermore, we attempted to use 3D-structure prediction to compare the strength of protein-DNA interactions between these replisome proteins and DNA with varying G/C-contents, but this proved uninformative because none of the interactions gave high enough interface predicted template modelling confidence scores to draw any reliable conclusion. Therefore, our modelling efforts did not reveal any clear structural differences between the replisomes of *P. falciparum, P. knowlesi* and *H. sapiens*.

## DISCUSSION

We present here the first direct comparison of replication dynamics in two ostensibly similar genomes with dramatically different nucleotide compositions. We conclude that the unusual features seen specifically in *P. falciparum* replication have evolved alongside, or owing to, this parasite’s extremely biased genome, and are not general to *Plasmodium* parasites. They are not, for example, imposed by the unusual replicative mode of schizogony, by auxotrophy for pyrimidines, or by some other factor imposed by replicating inside erythrocytes.

At the cellular level, we confirm that the dynamics of ORC deposition are similar in *P. knowlesi* and *P. falciparum* (8) (Figure 3), with ORC being synthesised and deposited prior to the start of S-phase, and not apparently degraded or exported after each replicative round and each karyokinesis. Thus, re-replication is evidently not prevented, in this syncytial system, by deactivation of ORC, as it is in human cells (32). Nevertheless, the differing genome compositions of *P. knowlesi* and *P. falciparum* evidently do impose different conditions on the deposition of ORC. We previously showed in *P. falciparum* that ORC bound preferentially to relatively high-G/C sequences, meaning that it tended to be found in CDS (the highest-G/C regions of a very A/T-biased genome) (8). Here, the equivalent tagged ORC1 protein in *P. knowlesi* did not show any significant enrichment in ChIP (data not shown) – possibly because ORC is entirely sequence-agnostic in a balanced genome, whereas the only factor generating clear ORC peaks in *P. falciparum* was the preference for G/C content higher than the low genomic average. Nevertheless, we could still show that regions mapped via DNAscent as active origins in *P. knowlesi* were not conspicuously G/C-rich relative to the *P. knowlesi* genomic average, so the balanced composition of this genome presumably makes most regions equally favourable for ORC binding.

Schizogony at the cellular level may look superficially similar in *P. knowlesi* and *P. falciparum*, but the dynamics of DNA replication at the molecular level differ greatly. Replication forks move at least ∼50% faster through *P. knowlesi* DNA than *P. falciparum* DNA (Figure 5). We previously saw this as a trend on DNA fibres (an orthogonal method, but challenging to use in this cell system) (2); here it was clearly quantified via DNAscent. Furthermore, replication fork speed increased slightly as S-phase progressed. The same has been reported in human cells (18,33): fork speeds are limited partly by nucleotide pools and these pools are ramped up during S-phase (11,12). Notably, however, we had previously recorded the opposite trend using two orthogonal methods in *P. falciparum,* with fork speeds diminishing across S-phase (8,10). At first, we hypothesised that nucleotide pools might simply be unable to keep up with the demands of replicating ten or more nuclei simultaneously at the end of schizogony, versus only one at the start (10), but these new data from *P. knowlesi* cast that hypothesis in doubt. Although nucleotide demands are likely less severe in *P. knowlesi* than *P. falciparum*, because *P. knowlesi* produces fewer daughter nuclei, and on average replicates slightly fewer of them at once (2), both species still replicate increasing amounts of DNA as schizogony progresses. In fact, the rise in actively replicating DNA seen in *P. knowlesi* (∼7-fold) was higher than the rise previously seen in *P. falciparum* (∼4-fold) (8). Therefore, *P. falciparum* DNA may simply be inherently more stressful for replication because of the high density of A/T mono- and di-nucleotide repeats, which will be prone to slippage and hairpin formation. Hence replication forks in *P. falciparum* are very slow overall, and perhaps become slower as schizogony proceeds because errors and stalls accumulate during this inherently stressed replication.

Another peculiarity of *P. falciparum* replication, not seen in *P. knowlesi*, was the strong bias to replicate faster in transcriptionally silent genes (8) (Figure 6). Again, because this is evidently not a general feature of small, gene-dense *Plasmodium* genomes, it may be imposed by genome bias. We speculate that slipped-strand-pairing, hairpins, etc., together with R-loops in active genes, may pose particular problems for genome replication in *P. falciparum* when replicative and transcriptional polymerases collide. Consistent with this, we saw in *P. falciparum* that the *var* gene family, which is generally silenced and also quite G/C-rich, was favoured for origin activation (8). The equivalent *SICAvar* family in *P. knowlesi* is also generally silenced, but was not apparently favoured.

Finally and most strikingly, in balanced genomes like *P. knowlesi* or human cells, DNA with very high A/T-content seemed to impose slowing on replication forks. By contrast, this was not seen in *P. falciparum*, where forks moved at their average speed even in DNA of >90% A/T (which is much higher than the genome-wide average). This suggests that the *P. falciparum* replisome may have evolved to tolerate its extreme genome bias, but at the expense of generally slow replication fork movement. We hypothesised that perhaps a polymerase or a secondary-structure helicase in *P. falciparum* might have an unusual structure, allowing it to associate more tightly with A/T-rich DNA, but neither 2D alignments nor 3D modelling revealed any insights on this hypothesis. In fact, the picture may be much more complex, placing it beyond the scope of this study.

*P. falciparum* is unique, thus far, in the detailed study of replication dynamics carried out by us (8,10) and other authors (34). However, it would be interesting to examine the equivalent dynamics in other heavily biased genomes, such as that of *D. discoideum* (77.6% A/T) (35). It would also be interesting (albeit impractical with current technology), to assemble a *P. falciparum* replisome *in vitro* and measure its dynamics through DNA sequences of different compositions. Finally, besides the academic interest of this study, the work could also inform the development of novel antimalarial drugs. Several such drugs, both historical and current, can damage or stall DNA replication (36) and we predict that synergistic combinations of such drugs could be particularly effective on the inherently slow and stressed replication of *P. falciparum* and other A/T-biased *Plasmodium* species.

## SUPPLEMENTARY FIGURE AND TABLE LEGENDS

Supplementary Figure 1. Plasmid maps used for tagging *P. knowlesi orc1* gene and subsequent genotyping and confirmation of successful tagging. (A) Cas9 plasmid with sgRNA targeting *P. knowlesi* A1-H.1 *orc1* (PKA1H_130007800); (B) Vector containing the donor DNA with ∼800 bp homology arms flanking the 3xHA tag followed by a skip peptide (T2A) and neomycin resistance (NeoR); (C) Genotyping of transfectants before (-) G418 and after (+) G418 selection using wild-type (WT) and mutant (M) directed primers; WT primers will give 917 bp and 1876 bp fragments for wild-type and mutant parasites, respectively; M primers will give a 1009 bp fragment in mutant parasites and none for wild-type parasites; (D) Western blot of fractionated protein lysates showing a majority of HA-tagged protein in the nuclear soluble fraction (NS). GAPDH and histone H4 are shown as control, present solely in the cytosolic (C) and insoluble nuclear fraction (NI), respectively. We noted a similar but more pronounced double banding of the probed tagged protein compared to what we have observed *P. falciparum* (8). This could be a result of a small N-terminal truncation, thus producing a slightly smaller band.

Supplementary Figure 2. Chromosomal distribution of DNAscent replication forks and origins in relation to *SICAvar*/*var* genes. Chromosome plots showing replication forks and origins in relation to *SICAvar* genes (A) and fork density plots over *SICAvar* genes during mid-schizogony (B) in *P. knowlesi*. Similar plot showing distribution of replication forks and origins (C) and fork density over *var* genes in *P. falciparum* (D). On the chromosome plots, black bars represent *SICAvar*/*var* gene locations, blue bars represent replication origins during early schizogony origins, and green bars represent origins at mid-schizogony. Light blue and dark blue line plots above the chromosomes represent replication fork density during early and mid-schizogony, respectively. Light green and dark green line plots below the chromosomes represent replication fork density during early and mid-schizogony, respectively. Fork density plots show mean fork density over *SICAvar*/*var* genes ± 1 kb, and significant difference was calculated to compare fork density at the start versus at the end of the genes.

Supplementary Table 1. List of oligonucleotide and primer sequences used for tagging *P. knowlesi orc1* gene and subsequent genotyping. Sequences of primers used to amplify fragments to construct the donor DNA, guide RNA oligos, sequencing and genotyping primers.

Supplementary Table 2. Summary of the total number of replication forks and origins called by DNAscent in *P. falciparum* (8) and *P. knowlesi*.

Supplementary Table 3. Genome-wide correlation between the localisation of HP1 and DNAscent forks and origins.

Supplementary Table 4. Percent identity matrices between human, *P. knowlesi* and *P. falciparum* polymerases (α, δ, and ε) and helicases (RECQ1 and WRN).

## ACKNOWLEDGEMENTS

We would like to thank Dr. Robert Moon and Dr. Franziska Mohring for the original CRISPR-Cas9 plasmids and for their advice on the transfection protocol. We would also like to acknowledge Dr. Richard Bartfai and Jonas Gockel (Radboud University) for their help in the *P. knowlesi* ORC1 ChIP-seq, and the lab of Prof. Julian Rayner for the thymidine kinase-expressing *P. knowlesi* parasite line.

## DATA AVAILABILITY

Plasmids (p45_Cas9_PkOrc1_sgRNA, and PkOrc1_donor_DNA) were made from plasmids originally generated by Dr. Robert Moon. The transgenic parasite line (*P. knowlesi* ORC1-3xHA + pTK-Pyr) used in this study may be available upon request through the corresponding authors. DNAscent data have been deposited at GEO, under accession number to follow. DNAscent v3.1.2, as well as DNAscent detect, DNAscent forkSense, and dnascent2bedgraph which are all part of DNAscent v3.1.2, are available under GPL-3.0 at https://github.com/MBoemo/DNAscent (also available on Zenodo at https://doi.org/10.5281/zenodo.7590289). Any additional information required to reanalyse the data reported in this paper is available from the corresponding authors upon request.

