## Supplementary material for "DNA replication dynamics are associated with genome composition in *Plasmodium* species": Supp Figs 1-2, Tables 1-4

### Supplementary Figure 1

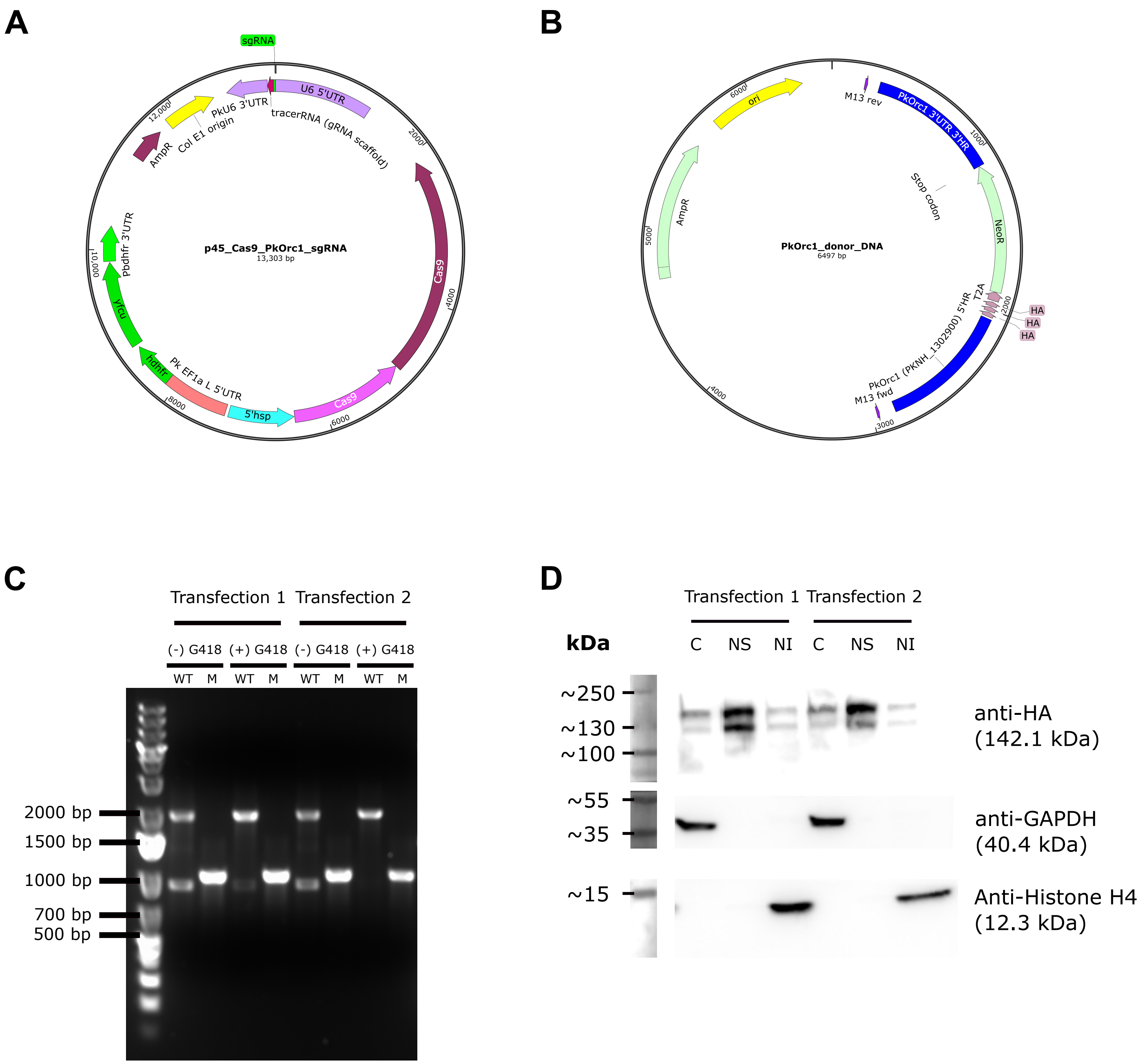

### Supplementary Figure 2

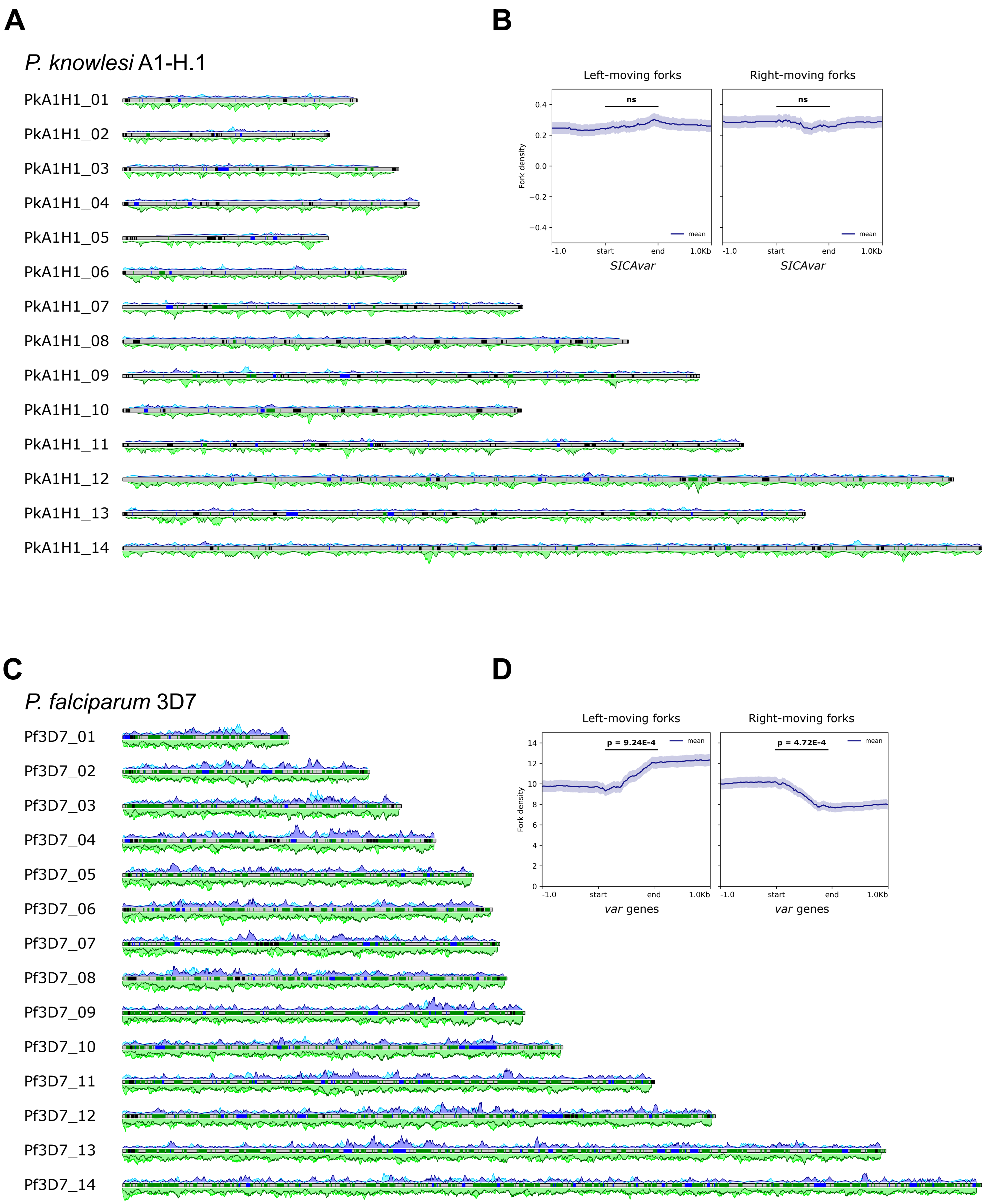

### Supplementary Table 1

| Primer name | Target | Details | Sequence | Fragment size (bp) |
|---|---|---|---|---|
| OH_FW (ol90) | Genomic DNA | Product A1: 5'Overhang-5'HR-HA_overlap | AATAGACGAGATTGATTACC | 893 |
| 5'_HR_RV (ol36) | | | CGTATGGGTACATGGTGGTACCGTAGAAGTTGAGCTTCTG | |
| HA_FW (ol54) | pSLI plasmid (*) | Product A2: 3xHA-T2A-NeoR-stop_codon | GGTACCACCATGTACCC | 963 |
| NeoR_RV-stop (ol55b) | | | TTAGAAGAACTCGTCAAG | |
| 3'_HR_NeoR_FW (ol37b) | Genomic DNA | Product A3: NeoR-stop_codon_overlap-3'HR-3'Overhang | CGCCTTCTTGACGAGTTCTTCTAACCATCTCTGCGGATAAAC | 909 |
| OH_RV (ol97) | | | GAATAATGGGAAGGATGG | |
| 5'_HR_FW (ol35) | Fused product A1:A2 and Product A3 | Product: 5'HR-3xHA-T2A-NeoR-stop_codon-3'HR | ACTAAGATAAACAGTAAATTGG | 2565 |
| 3'_HR_RV (ol38) | | | GGGAACCCTTCTAACATG | |
| NEB_sgRNA1_FW | *P. knowlesi* A1H1 *orc1* (PKA1H_130007800) | sgRNA with 5' and 3' overhangs for NEBuilder assembly (positive strand) | CCATATATTCTGAGTTACAGTATATTATT**ACTTCTACTAGCCATCTCTG**GTTTTAGAGCTAGAA | 72 |
| NEB_sgRNA1_RV | | sgRNA with 5' and 3' overhangs for NEBuilder assembly (negative strand) | CTTGCTATTTCTAGCTCTAAAAC**CAGAGATGGCTAGTAGAAGT**AATAATATACTGTAACTCAG | |
| M13 forward | Donor DNA sequencing | to confirm donor DNA sequence | GTAAAACGACGGCCAG | |
| M13 reverse | | | CAGGAAACAGCTATGAC | |
| HA1_sequencing | | | CGATGTTCCAGATTACGC | |
| NeoR_sequencing | | | CTTCTAACCATCTCTGCG | |
| sgRNA_sequencing | guide RNA sequencing | to confirm successful guide RNA insertion into Cas9 plasmid | CATTGTTCCCCCCTTTGTTTTGCAAG | |
| GT_PkOrc1_FW | Mutant specific | | AATAGACGAGATTGATTACC | 1009 |
| GT_T2A_RV | | | TGTTAATAAACTTCCTCTTCC | |
| GT_PkOrc1_3UTR_RV | WT specific | Used with GT_PkOrc1_FW forward primer | CTGCTTCGAAATTATTAGC | 917 (WT)<br>1876 (Mutant) |
| *pSLI-Orc1-3xHA used in Totanes, et al. Nucleic Acid Research (2023) | | | | |

### Supplementary Table 2

| | P. falciparum | | P. knowlesi | |
|---|---|---|---|---|
| | 30 hpi | 36 hpi | 20 hpi | 23 hpi |
| Left forks | 882 | 13021 | 815 | 997 |
| Right forks | 867 | 12877 | 787 | 958 |
| Origins | 228 | 3131 | 154 | 181 |

### Supplementary Table 3

| Correlation with HP1 | | Pearson R |
|---|---|---|
| *P. knowlesi* | 20hpi origins | -0.067 |
| | 20hpi leftForks | -0.1543 |
| | 20hpi rightForks | -0.1021 |
| | 23hpi origins | -0.0625 |
| | 23hpi leftForks | -0.1663 |
| | 23hpi rightForks | -0.1073 |
| *P. falciparum* | 30hpi origins | -0.0069 |
| | 30hpi leftForks | -0.0709 |
| | 30hpi rightForks | -0.0641 |
| | 36hpi origins | 0.0727 |
| | 36hpi leftForks | -0.0375 |
| | 36hpi rightForks | -0.0717 |

### Supplementary Table 4

**Polymerase α**

| | Plasmodium knowlesi DNA pol1 putative | Plasmodium falciparum DNA pol α subunit B | Homo sapiens pol α | Plasmodium knowlesi DNA pol α | Plasmodium falciparum DNA pol α |
|---|---|---|---|---|---|
| Plasmodium knowlesi DNA pol1 putative | 100.00 | 19.94 | 13.08 | 18.87 | 21.00 |
| Plasmodium falciparum DNA pol α subunit B | 19.94 | 100.00 | 16.22 | 18.89 | 21.61 |
| Homo sapiens pol α | 13.08 | 16.22 | 100.00 | 28.15 | 26.69 |
| Plasmodium knowlesi DNA pol α | 18.87 | 18.89 | 28.15 | 100.00 | 61.72 |
| Plasmodium falciparum DNA pol α | 21.00 | 21.61 | 26.69 | 61.72 | 100.00 |

**Polymerase δ**

| | Homo sapiens pol δ subunit A | Plasmodium falciparum pol δ subunit A | Plasmodium knowlesi pol δ subunit A | Homo sapiens pol δ small subunit | Plasmodium falciparum pol δ small subunit | Plasmodium knowlesi pol δ small subunit |
|---|---|---|---|---|---|---|
| Homo sapiens pol δ subunit A | 100.00 | 43.76 | 43.25 | 18.16 | 14.16 | 14.24 |
| Plasmodium falciparum pol δ subunit A | 43.76 | 100.00 | 84.00 | 16.48 | 21.28 | 20.35 |
| Plasmodium knowlesi pol δ subunit A | 43.25 | 84.00 | 100.00 | 15.89 | 19.83 | 18.82 |
| Homo sapiens pol δ small subunit | 18.16 | 16.48 | 15.89 | 100.00 | 30.86 | 34.52 |
| Plasmodium falciparum pol δ small subunit | 14.16 | 21.28 | 19.83 | 30.86 | 100.00 | 58.53 |
| Plasmodium knowlesi pol δ small subunit | 14.24 | 20.35 | 18.82 | 34.52 | 58.53 | 100.00 |

**Polymerase ε**

| | Homo sapiens pol ε subunit B | Plasmodium falciparum pol ε subunit B | Plasmodium knowlesi pol ε subunit B | Homo sapiens pol ε subunit A | Plasmodium falciparum pol ε subunit A | Plasmodium knowlesi pol ε subunit A |
|---|---|---|---|---|---|---|
| Homo sapiens pol ε subunit B | 100.00 | 19.77 | 19.49 | 14.53 | 17.77 | 16.54 |
| Plasmodium falciparum pol ε subunit B | 19.77 | 100.00 | 75.96 | 16.75 | 22.50 | 19.63 |
| Plasmodium knowlesi pol ε subunit B | 19.49 | 75.96 | 100.00 | 16.75 | 22.50 | 18.39 |
| Homo sapiens pol ε subunit A | 14.53 | 16.75 | 16.75 | 100.00 | 30.45 | 30.53 |
| Plasmodium falciparum pol ε subunit A | 17.77 | 22.50 | 22.50 | 30.45 | 100.00 | 75.36 |
| Plasmodium knowlesi pol ε subunit A | 16.54 | 19.63 | 18.39 | 30.53 | 75.36 | 100.00 |

**RECQ1**

| | Homo sapiens RECQ1 | Plasmodium falciparum RECQ1 | Plasmodium knowlesi RECQ1 |
|---|---|---|---|
| Homo sapiens RECQ1 | 100.00 | 29.87 | 31.59 |
| Plasmodium falciparum RECQ1 | 29.87 | 100.00 | 60.94 |
| Plasmodium knowlesi RECQ1 | 31.59 | 60.94 | 100.00 |

**WRM**

| | Homo sapiens WRN | Plasmodium falciparum WRN | Plasmodium knowlesi WRN |
|---|---|---|---|
| Homo sapiens WRN | 100.00 | 29.00 | 30.83 |
| Plasmodium falciparum WRN | 29.00 | 100.00 | 53.78 |
| Plasmodium knowlesi WRN | 30.83 | 53.78 | 100.00 |
